# Mutations in the zebrafish *hmgcs1* gene reveal a novel function for isoprenoids during red blood cell development

**DOI:** 10.1101/508531

**Authors:** Jose A. Hernandez, Victoria L. Castro, Nayeli Reyes-Nava, Laura P. Montes, Anita M. Quintana

## Abstract

Erythropoiesis is the process by which new red blood cells (RBCs) are formed and defects in this process can lead to anemia or thalassemia. The GATA1 transcription factor is an established mediator of RBC development. However, the upstream mechanisms that regulate the expression of *GATA1* are not completely characterized. Cholesterol is one potential upstream mediator of *GATA1* expression because previously published studies suggest that defects in cholesterol synthesis disrupt RBC differentiation. Here we characterize RBC development in a zebrafish harboring a single missense mutation in the *hmgcs1* gene (Vu57 allele). *hmgcs1* encodes the first enzyme in the cholesterol synthesis pathway and mutation of *hmgcs1* inhibits cholesterol synthesis. We analyzed the number of RBCs in *hmgcs1* mutants and their wildtype siblings. Mutation of *hmgcs1* resulted in a decrease in the number of mature RBCs, which coincides with reduced *gata1a* expression. We combined these experiments with pharmacological inhibition and confirmed that cholesterol and isoprenoid synthesis are essential for RBC differentiation, but that *gata1a* expression is isoprenoid dependent. Collectively, our results reveal two novel upstream regulators of RBC development and suggest that appropriate cholesterol homeostasis is critical for primitive erythropoiesis.

**Key Points:**

1. The products of the cholesterol synthesis pathway regulate red blood cell development during primitive erythropoiesis.
2. Isoprenoids regulate erythropoiesis by modulating the expression of the GATA1 transcription factor.

## Introduction

Erythropoiesis is the process of producing and replenishing the number of circulating red blood cells (RBCs). There are two unique waves of erythropoiesis: the primitive wave and the definitive wave. Erythropoiesis is tightly controlled and regulated by a balance of cell proliferation, differentiation, and survival ^1, 2^. The overproduction of RBCs or lack of RBCs can cause human disease. Diamond-Blackfan anemia and sickle cell anemia are two examples of rare congenital anomalies that arise from defects in the production of RBCs^3^, and polycythemia occurs as a consequence of too many RBCs ^4–6^. Genetic disorders of RBCs have revealed critical mediators of erythropoiesis ^7–11^, many of which include transcription factors. For example, Diamond-Blackfan anemia can result from mutations in the transcription factor *GATA1* ^12–14^. GATA1 is the founding member of the GATA family of zinc finger transcription factors ^15^ and interacts with a multitude of other proteins such as Friend of GATA (FOG), EKLF, SP1, p300 and PU.1 to promote erythropoiesis ^16^.

Cholesterol is one known regulator of RBC function as it maintains the structure and integrity of the RBC membrane and aids in the protection against oxidative stress^17–22^. But both *in vitro* and *in vivo* studies have raised the possibility that cholesterol biosynthesis regulates the differentiation of RBCs ^23, 24^. Knockdown of *OSC/LSS*, which catalyzes the cyclization of monoepoxysqualene to lanosterol, decreased the self-renewing capacity of K562 cells *in vitro* and results in increased cell death of progenitor like cells. Follow up *in vivo* assays have reinforced this premise as reduced cholesterol synthesis was associated with deficits in terminal RBC development ^24^. These data provide strong evidence that cholesterol’s function in RBCs is not restricted to membrane fluidity.

The cholesterol synthesis pathway (CSP) begins with synthesis of HMG-CoA from aceto-acetyl-CoA, which then undergoes several transformations to produce farnesyl pyrophosphate (FPP). FPP represents a branch point in the pathway, ultimately resulting in the production of cholesterol or isoprenoids ^25–27^. Both classes of lipids have diverse functions spanning membrane fluidity, protein prenylation, and precursors to various different types of molecules including vitamin D3. Cholesterol homeostasis has been previously linked to hematopoietic stem cell (HSC) differentiation ^28, 29^ and we confirmed that cholesterol synthesis is essential for RBC development ^24^. The function of isoprenoids is less clear because isoprenoids give rise to diverse molecules which themselves are critical for cell differentiation ^30–33^.

Here we show that the products of the CSP are essential for RBC development. We show that defects in cholesterol and/or isoprenoids results in deficient numbers of RBCs, but that each lipid regulates RBC number by unique mechanisms. We show that inhibition of isoprenoid synthesis disrupts the number of Gata1 positive cells produced, but the inhibition of cholesterol has no effect on *gata1* expression or the number of Gata1 positive cells. Thus, we demonstrate an essential function for the CSP during RBC specification and primitive erythropoiesis.

## Methods

### Zebrafish Care

For all experiments, embryos were obtained by crossing AB wildtype, Tupfel Long Fin wildtype, or *hmgcs1*^Vu57 34^. All embryos were maintained in embryo medium at 28° C and all experiments were performed according to protocol 811689-5 approved by The University of Texas El Paso Institutional Animal Care and Use Committee (IACUC). Genotyping was performed as previously described ^35^.

### Drug treatments and morpholino injection

Atorvastatin (Sigma, pharmaceutical grade, St. Louis MO), lonafarnib (Sigma, St. Louis MO), and Ro 48 8071 (Santa Cruz Biotechnology, Santa Cruz, CA) were each dissolved in 100% dimethyl sulfoxide (DMSO). Treatment was initiated at the sphere developmental stage (approximately 4-5hpf) and fresh drug was added every 18-24 hours until the harvest time points indicated in the figure legends. Drug concentrations were determined using a gradient of each drug (Supplementary Figure 1) and the concentration selected was based upon working conditions from previous literature. We selected a maximum tolerated sub-lethal dose producing a consistent phenotype according to a Fisher’s exact T-test as previously described in Quintana et. al. 2017 ^35^. Drugs were diluted in embryo medium to make working solutions at the following concentrations: 2.0 uM atorvastatin ^34–36^, 8 uM lonafarnib ^34, 37^, and 1.5uM Ro 48 8071. Final concentration of DMSO was less than 0.01 % in all samples and vehicle control treatment. Ro 48 8071 specificity for oxido-squalene synthesis has been previously described ^24, 24, 38, 39^. The specificity of lonafarnib has been previously described as a farnyesl protein transferase inhibitor ^40^. For morpholino injections, antisense *hmgcs1* morpholinos (AATCATATAACGGTGTTGGTTCGTG) were injected (0.025mM) at the single cell stage and fixed at the indicated time points within the figure legend. For all treatment groups (drug treatment and morpholino) statistical significance was obtained using a Fisher’s Exact Test.

### *o*-dianisidine staining

*o*-dianisidine (Sigma, St. Louis, MO) staining was performed as previously described by Paffett-Lugassy and Zon ^41^. Briefly, embryos were harvested at the desired time point and stained in the dark for 15 minutes at room temperature with *o*-dianisidine (Alfa Aesar, MA) (0.6mg/mL), 0.01M sodium acetate (Fisher, MA), 0.65% H_2_O_2_ (Fisher, MA), and 40% ethanol (Fisher, MA). Stained embryos were fixed with 4% paraformaldehyde (Electron Microscopy Sciences, PA) for 1 hour at room temperature and bleached using 3% hydrogen peroxide (H_2_O_2_) and 2% potassium hydroxide (KOH) (Fisher, MA) for 12 minutes. Embryos were washed with phosphate buffered saline (PBS) and stored in 4°C. Embryos were imaged with Zeiss Discovery Stereo Microscope fitted with Zen Software.

### Hemoglobin quantification

For hemoglobin quantification, larvae (numbers indicated in each figure graph) were homogenized with a pestle in purified water at 4 days post fertilization (dpf). Hemoglobin was measured with the Hemoglobin Assay Kit (Sigma-Aldrich, St. Louis, MO) according to manufacturer’s protocol. For analysis of the Vu57 allele, larvae were separated via distinct phenotypic hallmarks described previously ^34, 35^. The control contained both homozygous wildtype and heterozygous individuals harboring the Vu57 allele. Experiments were performed in a minimum of biological duplicate. For drug treatment assays, wildtype embryos were treated as described above before assaying for hemoglobin concentration. Statistical significance was determined using a T-test.

### Whole Mount In Situ Hybridization (WMISH) and Quantitative Real Time PCR (QPCR)

Whole mount *in situ* hybridization was performed as described by Thisse and Thisse ^42^. Briefly, embryos were harvested and dechorionated at the indicated time point and fixed in 4% paraformaldehyde (PFA) (Electron Microscopy Sciences, PA) 1 hour at room temperature (RT). Embryos were dehydrated using a methanol: PBS gradient and stored in 100% methanol overnight in −20°C. Embryos were rehydrated using PBS: Methanol gradient, washed in PBS with 0.1% Tween 20 and permeabilized with proteinase K (10ug/ml) for the time indicated by Thisse and Thisse ^42^. Permeabilized embryos were prehybridized in hybridization buffer (HB) (50% deionized formamide (Fisher, MA), 5X SSC (Fisher, MA), 0.1% Tween 20 (Fisher, MA), 50μg ml^-1^ heparin (Sigma, St. Louis, MO), 500μg ml^-1^ of RNase-free tRNA (Sigma, St. Louis, MO), 1M citric acid (Fisher, MA) (460μl for 50ml of HB)) for 2-4 hours and then incubated overnight in fresh HB with probe (*gata1a* 75ng, *hbbe1.1* 75ng, *alas2* 150ng) at 70°C. Samples were washed according to protocol, blocked in 2% sheep serum (Sigma, St. Louis, MO), 2 mg ml^-1^ bovine serum albumin (BSA) (Sigma, St. Louis, MO) for 2-4 hours at RT, and incubated with anti-DIG Fab fragments (1:10,000) (Sigma, St. Louis, MO) overnight at 4°C. Samples were developed with BM purple AP substrate (Sigma, St. Louis, MO) and images were collected with a Zeiss Discovery Stereo Microscope fitted with Zen Software. Statistical analysis was performed using a Fisher’s Exact Test. For QPCR, RNA was isolated from embryos at the indicated time point with Trizol (Fisher, MA) according to manufacturer’s protocol. Reverse transcription was performed using iScript (Bio-Rad, Redmond, WA) and total RNA was normalized across all samples. PCR was performed in technical triplicates for each sample using an Applied Biosystems StepOne Plus machine with Applied Biosystems associated software. Sybr green (Fisher, MA) based primer pairs for each gene analyzed are as follows: *gata1a* fwd GTTTACGGCCCTTCTCCACA, *gata1a* rev CACATTCACGAGCCTCAGGT, *hbbe1.1* fwd TGAATCCAGCACCCATCTGA, *hbbe1.1* rev CTCCGAGAAGCTCCACGTAG, *rpl13a* fwd TCCCAGCTGCTCTCAAGATT, *rpl13a* rev TTCTTGGAATAGCGCAGCTT. Analysis performed using 2^ΔΔct^. Statistical analysis of mRNA expression was performed using a T-test. All QPCR was performed in biological duplicate or triplicate.

### Confocal Imaging

Embryos were fixed at the given time point and then mounted in 0.6% low melt agar in a glass bottom dish (Fisher, MA). Imaging was performed on a Zeiss LSM 700 at 20X magnification. Images were restricted to the caudal hematopoietic tissue. For each fish, a minimum of 12-20 z-stacks were collected. Statistical significance was obtained using a T-test.

### Data Sharing

For access to datasets and protocols please contact corresponding author by email.

## Results

### Mutations in *hmgcs1* disrupt RBC development

Based upon previous data ^24^, we sought to determine the number of mature RBCs in a zebrafish harboring mutations in the *hmgcs1* gene (Vu57). The Vu57 allele introduces a single missense mutation (H189Q) in the *hmgcs1* gene, which encodes the first enzyme in the CSP. The VU57 allele abrogates cholesterol synthesis causing a multiple congenital anomaly syndrome characterized by defects in myelination, myelin gene expression, cardiac edema, pigment defects and craniofacial abnormalities ^34, 35^. We first detected the number of hemoglobinized RBCs in Vu57 homozygous mutants (*hmgcs1*^-/-^) or their wildtype siblings with *o*-dianisidine at 4 dpf. Over the first 4 days of development all of the circulating RBCs are derived from primitive erythropoiesis, therefore, analysis of hemoglobinzed RBCs at day 4 accurately depicts deficiencies in primitive erythropoiesis ^43^. Wildtype siblings had adequate numbers of hemoglobinized RBCs throughout development and at 4 dpf the RBCs lined the ventral head vessels of the neck and face (Figure 1A). Homozygous carriers of the Vu57 allele had a reduced number of cells populating the ventral head vessels. This decrease was not accompanied by a significant accumulation of cells in other regions of the body (Figure 1A’&B’).

**Figure 1:**
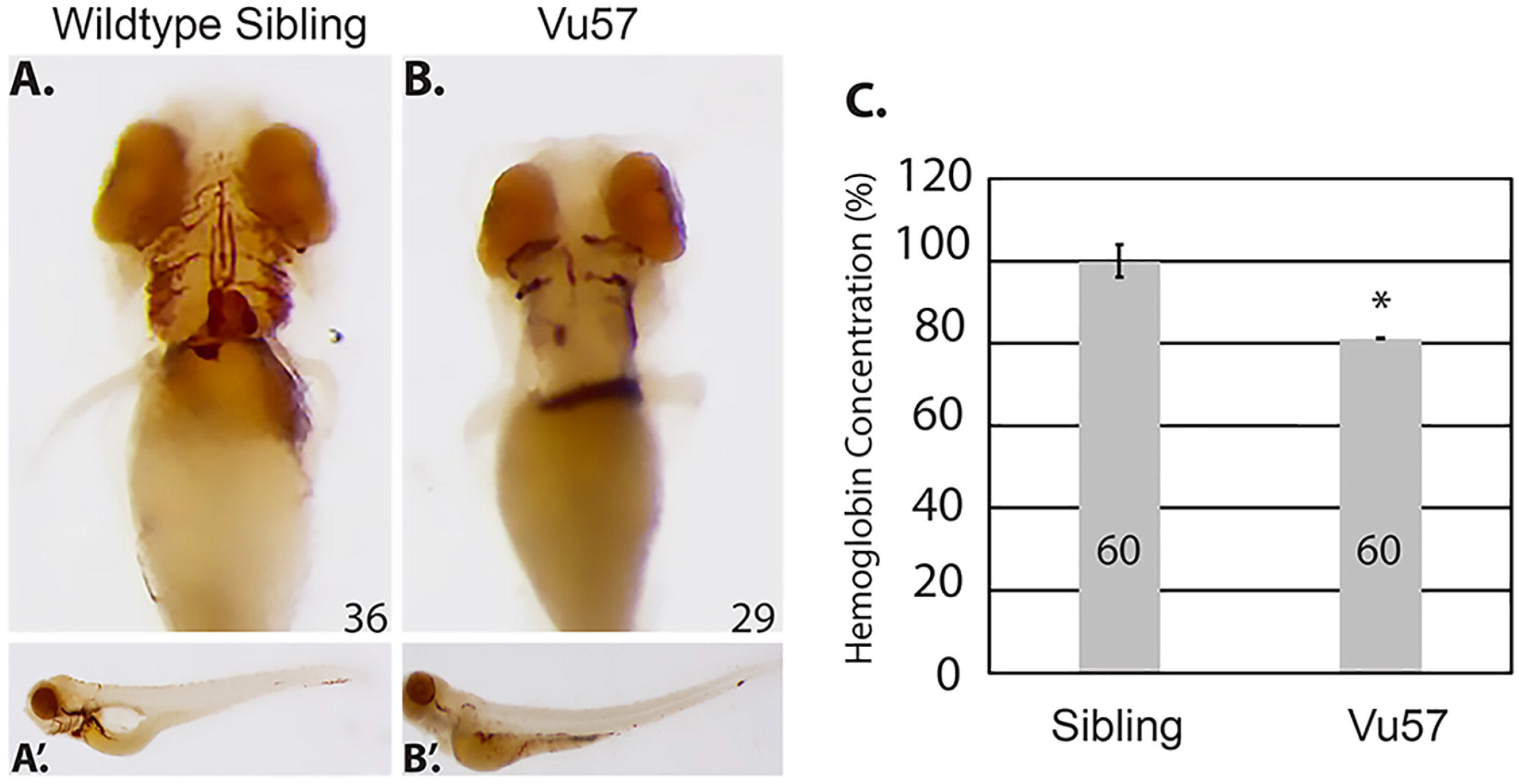
Mutations in *hmgcs1* cause a decrease in hemoglobinized red blood cells (RBCs). A-B *hmgcs1*^-/-^ (Vu57) and their wildtype siblings (*hmgcs1*^+/+^) were stained for hemoglobinized RBCs at 4 days post fertilization (n= 36 *hmgcs1*^+/+^, n=29 *hmgcs1*^-/-^) (dpf) using *o*-dianisidine. Ventral views of the hemoglobinized RBCs are shown in A&B, with full body images of both wildtype siblings and homozygous mutants (Vu57) shown in A’&B’. The phenotype was completely penetrant in homozygous mutants. Total numbers of animals were obtained across a minimum three biological replicates. C. The concentration of hemoglobin was measured in siblings (a pool of wildtype and heterozygous individuals) and embryos carrying the Vu57 allele. The assay was performed with two biological replicates with a total of 60 larvae.*p<0.05.

We next quantified the decrease in circulating RBCs in homozygous mutants and their siblings. We performed a quantitative measure of total hemoglobin content using a colorimetric assay in which endogenous hemoglobin can be measured quantitatively at a wavelength of 400nm ^44^. In order to measure the levels of hemoglobin in siblings and Vu57 carriers, homozygous Vu57 larval were separated according phenotype at 3 dpf ^34^ and the total hemoglobin content of homozygous carriers was compared to the hemoglobin content of wildtype and heterozygous siblings at 96 hpf. Phenotypic hallmarks of the Vu57 allele include craniofacial abnormalities and cardiac edema ^34^. As shown in Figure 1C, the total hemoglobin content in homozygous carriers of the Vu57 allele was decreased by approximately 20% (p<0.05). Taken together these data suggest that mutations in *hmgcs1* disrupt RBC development.

### Mutation of *hmgcs1* disrupts the expression of markers associated with RBC differentiation

One possible explanation for the loss of total hemoglobin content observed in larvae carrying the Vu57 allele could stem from an inability to produce globin mRNA. To determine whether mutations in *hmgcs1* interfere with globin expression, we performed whole mount *in situ* hybridization (ISH) at 26 hpf with an anti-hbbe1.1 riboprobe. *hbbe1.1* was expressed in the caudal intermediate cell mass (ICM) of wildtype siblings and the onset of circulation was readily apparent as *hbbe1.1* mRNA was detected over the yolk sac (Figure 2A). *hbbe1.1* expression was up-regulated in Vu57 embryos with expression that localized throughout the entire ICM and was not restricted to the most caudal region (Figure 2B arrowhead). In addition, *hbbe1.1* expression over the yolk sac was increased relative to wildtype siblings at 26 hpf (Figure 2A&B).

**Figure 2:**
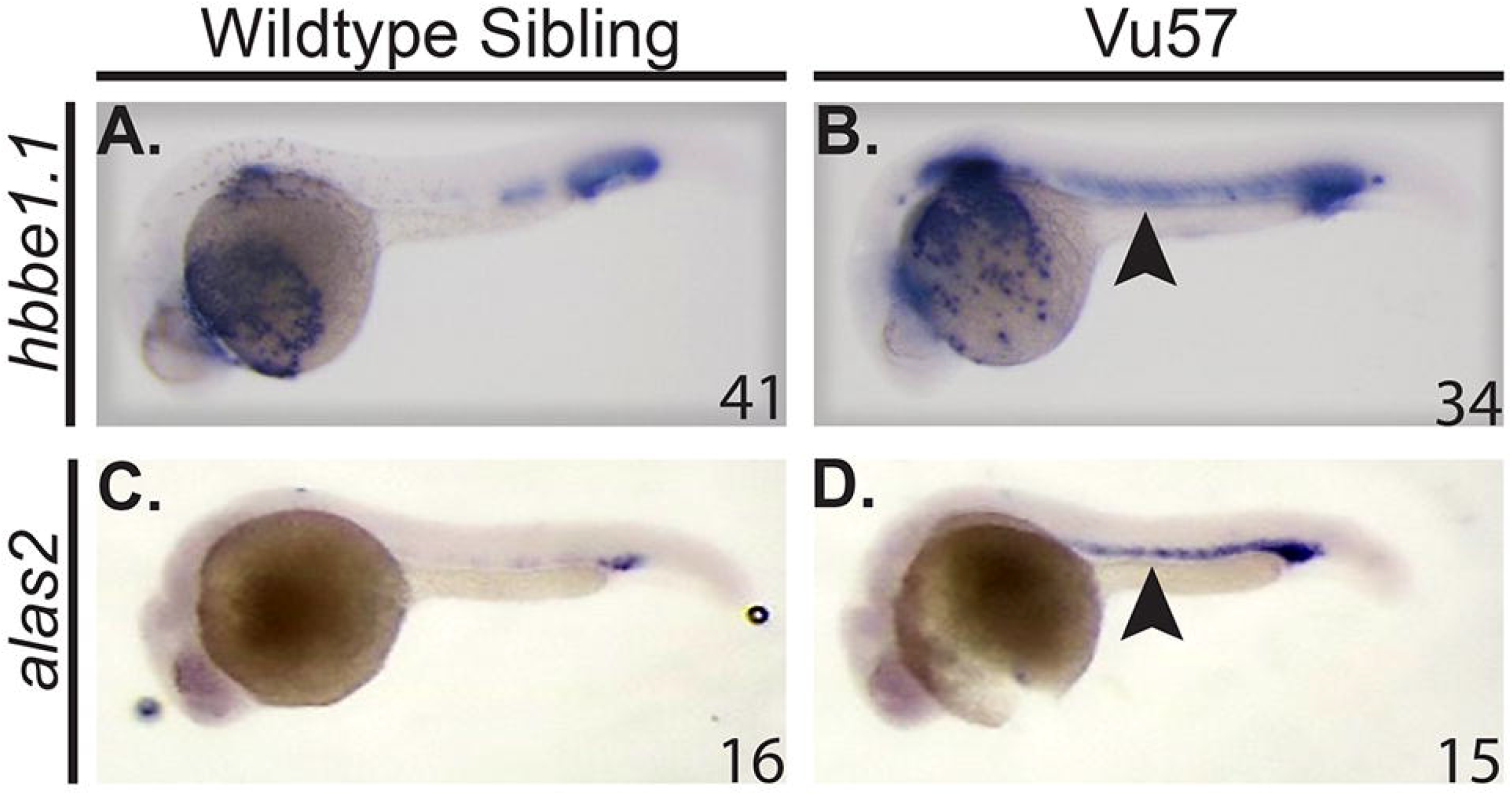
*hbbe1.1* and *alas2* are expressed in *hmgcs1* mutant larvae. A-D. Whole mount *in situ* hybridization (ISH) was performed to detect the expression of *hbbe1.1* (n= 41 *hmgcs1*^+/+^ (sibling), n=34 *hmgcs1*^-/-^(Vu57)), (A&B) or *alas2* (n= 16 *hmgcs1*^+/+^ (sibling), n=15 *hmgcs1*^-/-^ (Vu57)), (C&D) at 26 hours post fertilization. Purple indicates expression of each gene in the intermediate cell mass and areas of increased or abnormal expression are indicated by the arrowhead. Total number of animals was achieved with a minimum of two biological replicates.

We next measured the expression of *alas2*, which encodes the first enzyme in heme biosynthesis ^45^. Wildtype siblings expressed appropriate *alas2* expression in the caudal ICM at 26 hpf (Figure 2C), but the level of *alas2* in embryos with the Vu57 allele was spatially disrupted spanning the entire ICM (Figure 2D arrowhead) and increased relative to wildtype siblings. These data are consistent with the level and spatial expression of *hbbe1.1*, as *alas2* is known to modulate the levels of globin ^46^. Taken together, these data demonstrate that mutant embryos maintain the expression of globin and some of the enzymes necessary for heme synthesis.

### *gata1a* expression is decreased in *hmgcs1* mutant embryos

Given the abnormal expression of globin (*hbbe1.1*), we hypothesized that the mutation of *hmgcs1* disrupts the expression of *GATA1*, a known regulator of globin expression. We measured the expression of *gata1a*, the zebrafish ortholog of *GATA1* using ISH and quantitative PCR (QPCR) at 18 somites and 26 hpf in mutants and their wildtype siblings. At 18 somites, wildtype siblings expressed *gata1a* in the caudal ICM (Figure 3A, dorsal view), but the Vu57 allele resulted in decreased *gata1a* expression (Figure 3A-B arrowheads). This decrease in *gata1a* persisted through the onset of circulation, as we observed reduced *gata1a* expression at 26hpf (Figure 3C-D, arrowhead). We next quantified the expression of *gata1a* by QPCR. We quantified the expression of *gata1a* in embryos injected with an *hmgcs1* morpholino because genotyping of mutant larvae prior to RNA isolation could not be consistently achieved without rapid decay in total RNA quality. Microinjection of *hmgcs1* morpholinos accurately phenocopied the RBC deficits observed with the Vu57 allele (Supplementary Figure 2) and QPCR confirmed a near 70% reduction in *gata1a* expression in morphants (Figure 3A-D&3I, p<0.05).

**Figure 3:**
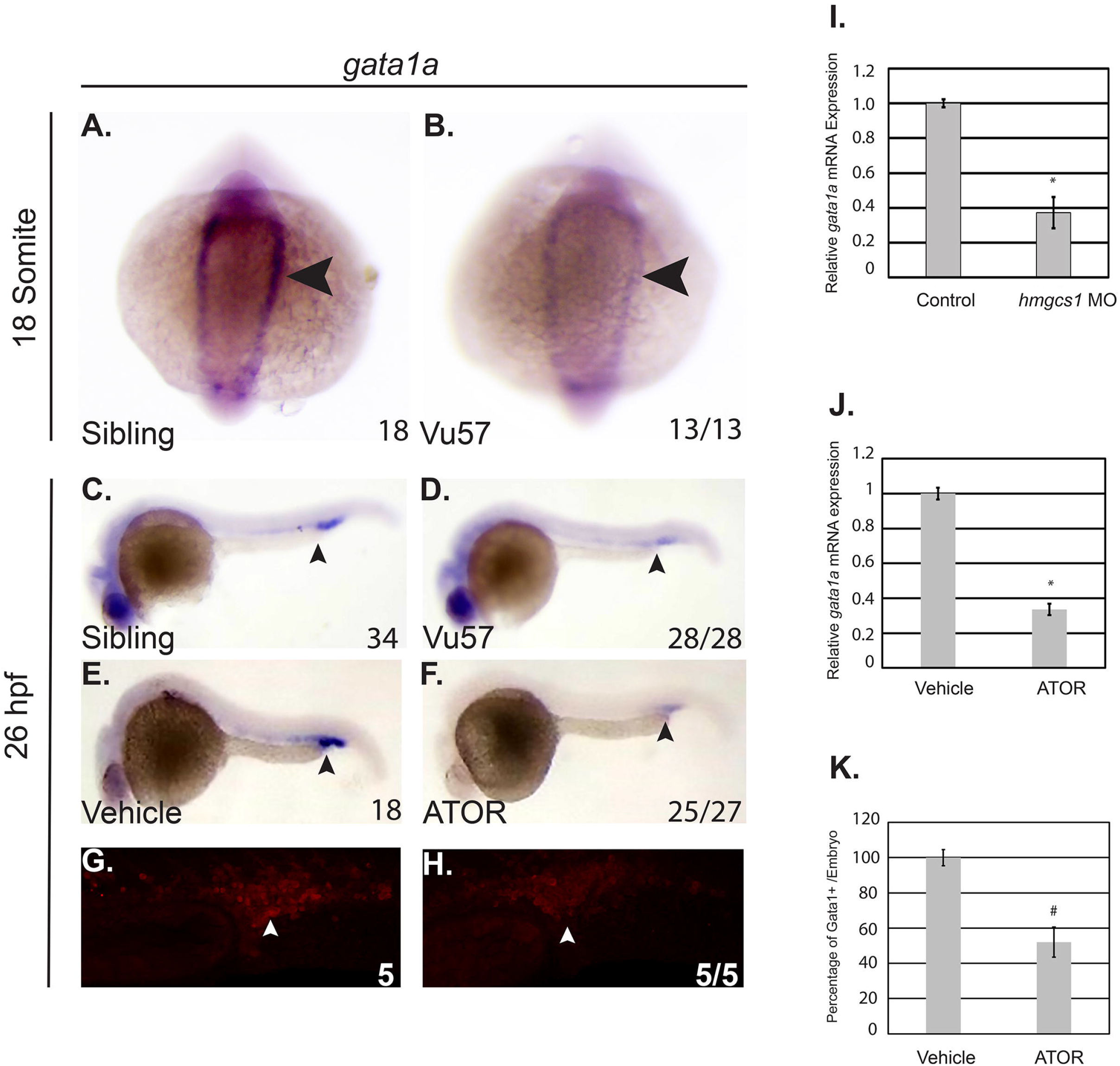
Mutation of *hmgcs1* disrupts *gata1a* expression. A-D. Whole mount *in situ* hybridization (ISH) was performed to detect the expression of the *gata1a* transcription factor at the 18 somite stage (n= 18 *hmgcs1*^+/+^ (Sibling), n=13 *hmgcs1*^-/-^ (Vu57)) and 26 hours post fertilization (hpf) (n= 34 Sibling, n=28 Vu57). Total numbers of animals were obtained with a minimum of two biological replicates. E&F. Embryos were treated with vehicle control (dimethyl sulfoxide (DMSO)) or 2uM atorvastatin (ATOR) (n= 18 DMSO, n=27 ATOR, p=0.0001) from sphere stage to 26 hpf and subjected to ISH to detect *gata1a* expression. Numbers of embryos affected are indicated below each figure. P-value represents a Fisher’s exact T-test demonstrating the numbers affected per treatment group. Total numbers of embryos were obtained across two biological replicates. Arrowheads indicate area of *gata1a* expression at each time point. G-H. *Tg*(*gata1a*:dsRed) embryos were treated with vehicle control (dimethyl sulfoxide (DMSO)) or 2uM atorvastatin. Fluorescence was visualized using a confocal microscope. The number of cells/Z-stack was quantified using ImageJ. I. Antisense *hmgcs1* morpholinos were injected (0.025mM) at the single cell stage and total RNA was extracted at the 18 somite stage. Quantitative real time PCR (QPCR) was performed to detect the expression of *gata1a*. All samples were performed in technical triplicate and error bars represent the standard deviation of technical triplicates. J. Total RNA was isolated from embryos treated with vehicle control or ATOR and QPCR was performed to detect the expression of *gata1a*. All samples were performed in technical triplicate and error bars represent the standard deviation of technical triplicates. *p<0.05 K. Quantification of the number of dsRed cells from G&H. (# p= 7.25273E-07).

We next measured *gata1a* expression in wildtype embryos treated with 2uM atorvastatin (ATOR), a drug that inhibits the rate-limiting step of the cholesterol synthesis pathway ^47^, and should mimic the effects of mutations in *hmgcs1. gata1a* expression was decreased in ATOR treated embryos relative to vehicle control (Figure 3E-F, arrowheads, p=0.0001) and QPCR confirmed that ATOR treatment caused a significant reduction in *gata1a* expression (Figure 3J, p<0.05). We next confirmed these results by treating *Tg*(*gata1a*:dsRed) larvae with 2uM ATOR or vehicle control. Treatment with ATOR caused an approximate 50% decrease in the number of dsRed positive cells (Figure 3G, H&K p=7.25273E-07). Collectively, these data suggest that the Vu57 allele decreases the number of Gata1a positive cells produced during primitive erythropoiesis.

### Cholesterol and Isoprenoids regulate RBC development

Mutation of *hmgcs1* disrupts the first enzyme of the CSP ^34^ effectively interfering with the production of both cholesterol and isoprenoids. Recent evidence suggests that each of these two lipids can regulate the same biological process, however by independent molecular and cellular mechanisms ^34, 35, 37^. We hypothesized that the defects observed in mutant larvae are cholesterol dependent. We treated wildtype embryos with either vehicle control (DMSO), 1.5uM Ro 48 8071, to inhibit cholesterol, but not isoprenoids, 8uM lonafarnib, to inhibit farnesylated isoprenoids, but not cholesterol or 2uM ATOR, a control to mimic the Vu57 allele. According to *o*-dianisidine, vehicle treated embryos (DMSO) exhibited the appropriate number and spatial organization of RBCs in the ventral head vessels at 4 dpf (Figure 4A). Notably, treatment with ATOR induced a cerebral hemorrhage that was not consistent with the Vu57 allele (Figure 4B). Embryos treated with 1.5uM Ro 48 8071 or 8uM lonafarnib had visibly fewer RBCs (Figure 4A-D p=0.0001), suggesting that cholesterol synthesis is required for RBC development. Cerebral hemorrhages were not observed upon treatment with Ro 48 8071 or lonafarnib. We further quantified the total hemoglobin content from larvae treated with each drug or vehicle control. As shown in Figure 4E, treatment with each drug resulted in a statistically significant decrease in total hemoglobin content. Drug treatment resulted in a more marked decrease in hemoglobin concentration relative to larvae harboring the Vu57 allele (Figure 1). This can likely be attributed to the fact that we performed a comparison between homozygous carriers of the Vu57 allele with a pool of heterozygous and wildtype homozygous individuals suggesting that heterozygous individuals demonstrate some degree of deficits in RBC development. Taken together, these data raise the possibility that the synthesis of cholesterol and isoprenoids is essential for RBC development/differentiation.

**Figure 4:**
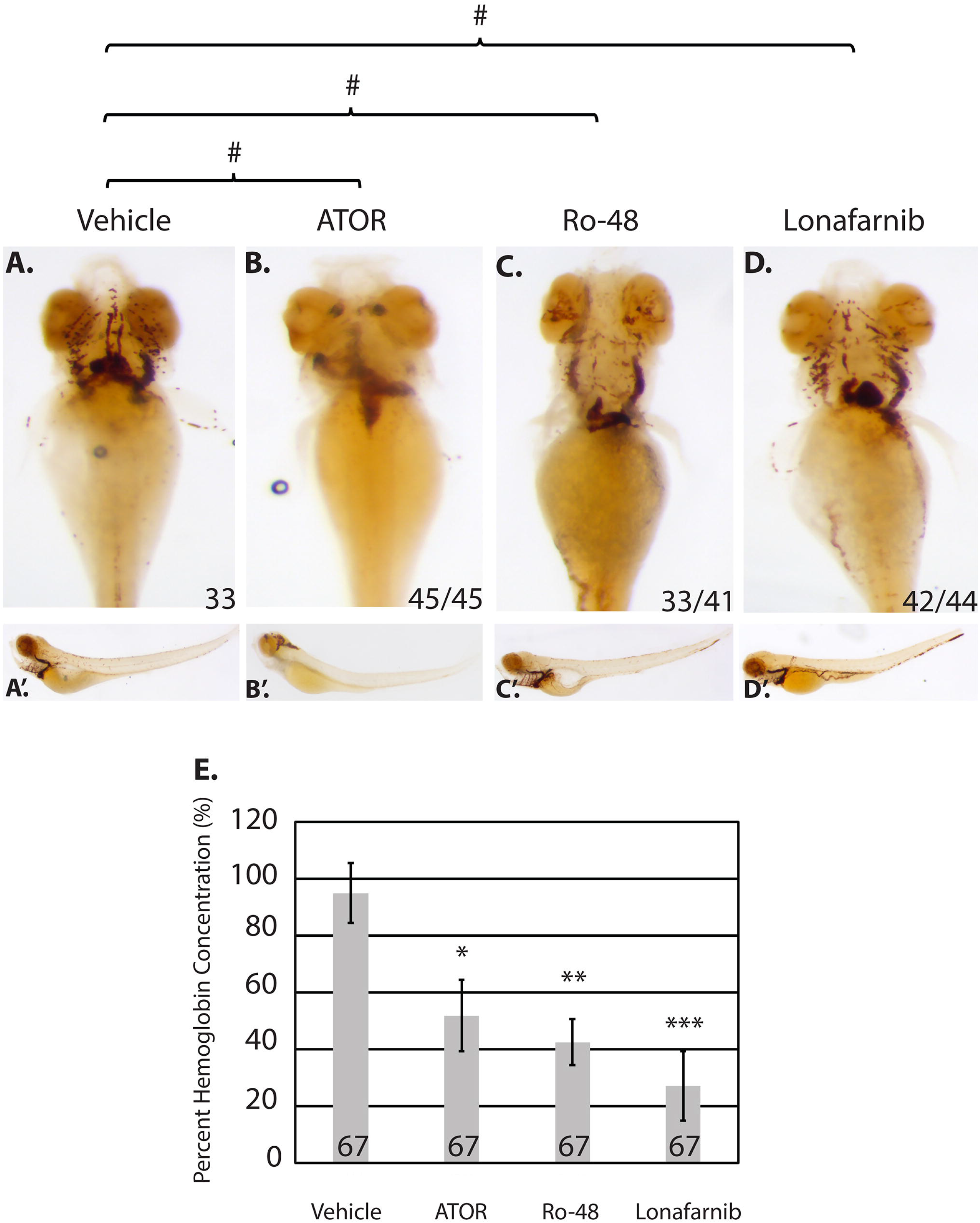
Cholesterol and isoprenoids regulate erythropoiesis. A-D. Embryos were treated at sphere stage with atorvastatin (ATOR), Ro 48 8071 (Ro-48) to inhibit cholesterol, lonafarnib to inhibit isoprenoids, or vehicle control (dimethyl sulfoxide). At 4 days post fertilization (dpf) embryos (n= 33 DMSO, n=45 ATOR n=41 Ro-48, and n=44 lonafarnib) were stained with *o*- dianisidine to observe mature red blood cells (RBCs). # p= 0.0001 p-value indicates the number of affected embryos affected is statistically significant according to Fisher’s Exact Test. (D) The concentration of hemoglobin was measured in embryos treated with atorvastatin (ATOR), Ro 48 8071 (Ro-48) to inhibit cholesterol, lonafarnib to inhibit isoprenoids, or vehicle control (dimethyl sulfoxide) at 4 days post fertilization. The number indicates total larvae analyzed across three biological replicates. *p= 0.000381218, **p= 2.20098E-05, ***p= 1.42E-05.

### *gata1a* expression is isoprenoid dependent

The Vu57 allele results in decreased *gata1a* expression and increased *hbbe1.1* expression. Therefore, we measured the expression of each gene in wildtype embryos treated with vehicle control (DMSO), 1.5uM Ro 48 8071, or 8uM lonafarnib using ISH at 26 hpf. *hbbe1.1* expression was localized to the caudal most region of the ICM and over the yolk sac in vehicle control embryos (Figure 5A). Inhibition of cholesterol or isoprenoids caused a statistically significant increase in *hbbe1.1* mRNA (Figure 5G, p<0.05) that was visible over the yolk sac (Figure 5A-C, p=0.0001). The level and spatial expression of *hbbe1.1* is consistent with those observed in embryos carrying the Vu57 allele (Figure 2). Interestingly, the expression of *gata1a* was not affected by treatment with 1.5uM Ro 48 8071, but was significantly decreased when isoprenoid synthesis was inhibited (Figure 5D-F; Figure 5H, p<0.05). The decrease in *gata1a* expression was consistent with a decrease in the number of Gata1^+^ cells as demonstrated by the *Tg*(*gata1a*:dsRed) (Figure 5I, p=2.7125E-07) Collectively, these data suggest that isoprenoids regulate RBC differentiation in a *gata1a* dependent manner.

**Figure 5:**
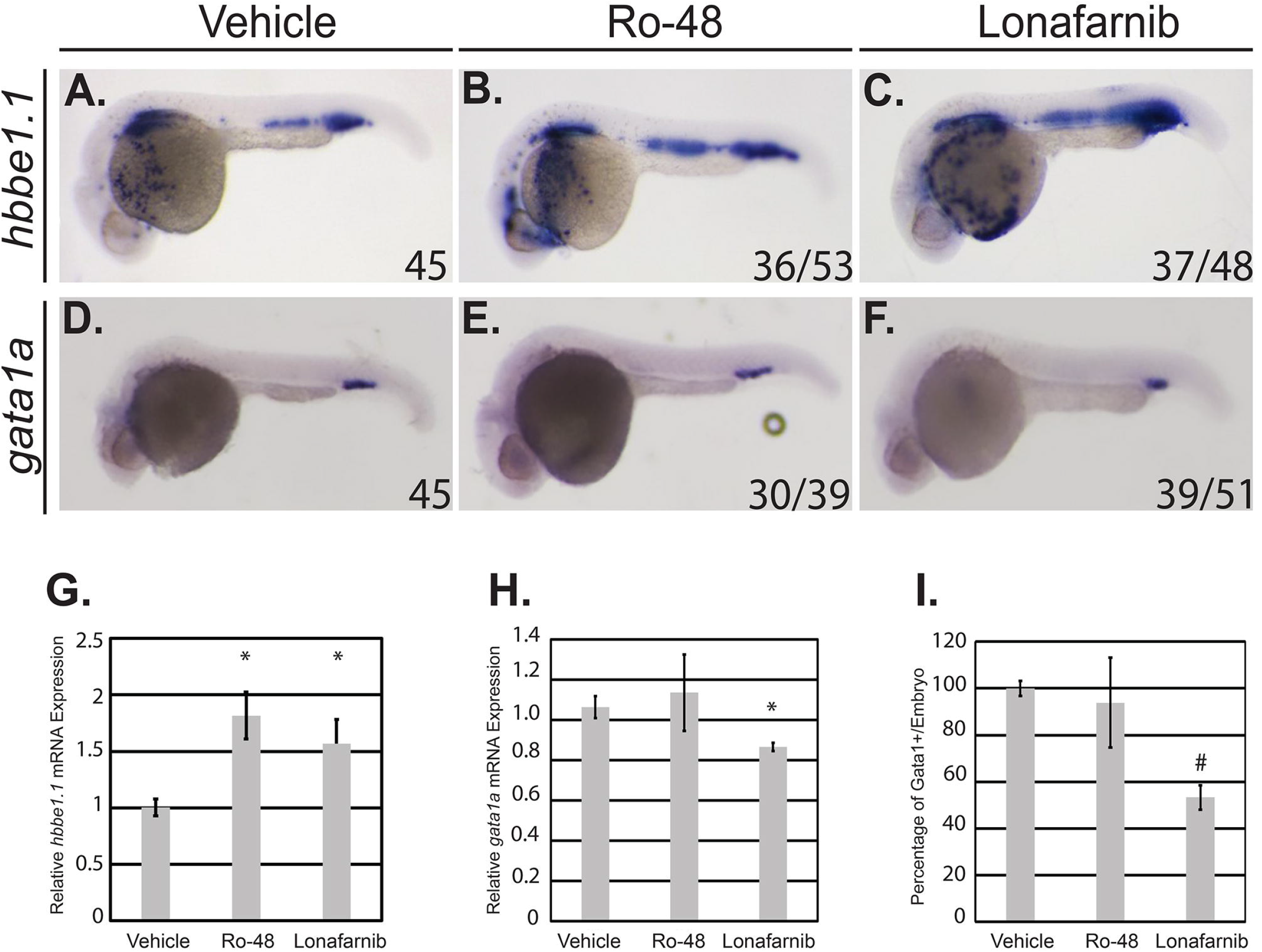
Isoprenoids regulate red blood cell (RBC) development in a *gata1a* dependent manner. A-F. Embryos were treated at sphere stage with Ro 48 8071 (Ro-48) to inhibit cholesterol, lonafarnib to inhibit isoprenoids, or vehicle control (dimethyl sulfoxide). At 26 hours post fertilization (hpf), whole mount *in situ* hybridization was performed to detect *hbbe1.1* (n= 45 DMSO, n=53 Ro-48 *(p=0.0001), and n=48 lonafarnib (p=0.0001)) (A-C) or *gata1a* (D-F) expression n= 45 DMSO, n=39 Ro-48 (NS) and n=51 lonafarnib, (p=0.0001)). Total embryos were obtained with a minimum 3 biological replicates. p-value designates statistical significance relative to vehicle control according to a Fisher’s exact test. G-H. Total RNA was isolated from embryos treated with vehicle control (dimethyl sulfoxide), Ro-48, or lonafarnib and QPCR was performed to detect the expression of *hbbe1.1* (G) or *gata1a* (H). All samples were performed in technical triplicate and error bars represent the standard deviation of technical triplicates. p<0.05. I. *Tg*(*gata1a*:dsRed) embryos were treated at sphere stage with Ro-48 to inhibit cholesterol, lonafarnib to inhibit isoprenoids, or vehicle control (dimethyl sulfoxide). Fluorescence was visualized using a confocal microscope. The number of cells/Z-stack was quantified using ImageJ. # p= 2.7125E-07.

## Discussion

Here we show that cholesterol and isoprenoids regulate erythropoiesis using a zebrafish harboring mutations in the *hmgcs1* gene (Vu57 allele). Mutations in human *HMGCS1* have not been associated with disease, but there are 8 congenital anomalies that occur as a consequence of mutations within other enzymes of the CSP ^5–11, 13, 48^. These congenital anomalies are characterized by diverse phenotypes ^7, 21, 48–50^ and mutations in the zebrafish *hmgcs1* gene mimics these disorders resulting in a multiple congenital anomaly syndrome. Therefore, zebrafish with mutations in *hmgcs1* have the potential to reveal novel cellular and molecular mechanisms underlying individual phenotypes across multiple genetic disorders.

Cholesterol represents approximately ½ the weight of a RBC membrane and governs membrane fluidity, transport, reversible deformations and survival in response to oxidative stress^51, 52^. In addition cholesterol is a precursor for multiple molecules including bile acid, vitamin D, and steroid hormones and deficiencies in cholesterol synthesis interfere with proper RBC development ^23, 24^. Based upon these data, we hypothesized that mutation of *hmgcs1* would disrupt erythropoiesis *in vivo*. We found that homozygous mutations in *hmgcs1* cause a decrease in the number of hemoglobinized RBCs and total hemoglobin content, consistent with previous work ^24^ demonstrating that defects in RBC number can be rescued by the exogenous injection of water soluble cholesterol. Our study using the Vu57 allele establishes that the products of the CSP are essential for proper RBC homeostasis, however, the mechanism by which the CSP exerts these effects is yet to be elucidated. Given the role of cholesterol in the RBC membrane, it is plausible that cholesterol regulates cell survival. However, the function of the CSP might not be limited to cell death or stress mechanisms because previously published work has indicated a regulatory function for cholesterol and its derivatives at the level of cellular differentiation ^53–60^. Future work that analyzes both primitive and definitive hematopoiesis at unique stages of differentiation in different model systems is likely to identify the exact cellular mechanisms underlying the phenotypic alteration we describe. The closely related zebrafish mutant of *hmgcrb* will be of great utility, as we hypothesize that mutation in *hmgcrb* ^36^ will produce overlapping phenotypes with the Vu57 allele.

The GATA family of transcription factors are essential mediators of erythropoiesis. Specifically, the expression of *GATA1* signals the commitment of a common myeloid progenitor towards an erythroid fate. Numerous studies have confirmed that the expression of *GATA1* is at the center of at least two axis governing cell fate decisions. The expression of GATA1 represses the expression of *GATA2*, a second member of the family whose expression promotes a progenitor cell fate^15, 61^. GATA1 expression also antagonizes the expression of *SPI-1*, which promotes myeloid differentiation ^62, 63^. Here we demonstrate for the first time that the expression of *gata1a*, the zebrafish ortholog of *GATA1*, is linked to the CSP. Moreover, we establish that expression of *gata1a* is isoprenoid dependent. These data are supported by previously published studies by Quintana et. al., which demonstrate that defects in cholesterol synthesis disrupt RBC differentiation without disrupting early *gata1a* expression ^24^. Despite reduced expression of *gata1a* in *hmgcs1* mutants, differentiating RBCs maintain their ability to initiate and maintain *globin* and *alas2* expression. The inhibition of the CSP did not cause accumulation of mature RBCs to other bodily regions except for in the presence of atorvastatin treatment, where cerebral hemorrhages are observed. This phenotype is not consistently observed with the Vu57 allele or larvae treated with lornafarnib or Ro 48 8071, but has been reported in *hmgcrb* mutants ^36^. Despite the presence of cerebral hemorrhages in atorvastatin treated embryos, we still observe a statistically significant decrease in total hemoglobin content in these larvae. Thus, suggesting that inhibition of the CSP reduces total hemoglobin content, which is consistent with an accumulation of *globin* and *alas2* RNA. These results are further supported by *in vitro* studies of *GATA1* deletion where *GATA1* negative cells undergo developmental arrest, but maintain expression of GATA target genes, including globin ^64^.

Our data establishes that isoprenoid synthesis is essential for appropriate *gata1a* expression. These effects are likely to be indirect because isoprenoids are a large class of molecules with diverse functions. Retinoids are an isoprenoids derivative ^65^ and retinoic acid is one potential regulator of blood cell differentiation because previous studies have established that retinoic acid signaling increases the number of HSCs ^30, 31, 33^ and mediates the formation of HSCs from the mesoderm ^66^. However, these mechanisms are likely to be complex and stage specific because retinoic acid has been shown to decrease the expression of *gata1* in zebrafish^67^. Thus, retinoic acid is only one potential mediator of erythropoiesis.

Here we demonstrate that cholesterol and isoprenoids, two products of the CSP modulate RBC differentiation *in vivo*. The cholesterol independent mechanisms disrupt *gata1a* expression and the number of Gata1a positive cells produced. This is notable as GATA1 regulates at least two axis regulating lineage fate decisions, but at the present time it is not clear if there are hematopoietic defects prior to onset of *gata1a* expression. The presence of cerebral hemorrhages in atorvastatin treated embryos may shed some light on this question, as HSCs and endothelial cell progenitors both arise from a common bipotent progenitor during primitive hematopoiesis ^68, 69^ and defects in both lineages could indicate early defects in the formation or differentiation of cells from mesoderm. Thus, future studies that define the mechanisms by which cholesterol and isoprenoids regulate all stages of differentiation, including early HSCs and myeloid cells, are warranted.

Our study focuses on the regulation of *GATA1* expression, primarily in the context of isoprenoids. Given the role of isoprenoids in development and signaling, it is likely that they are a positive upstream regulator of *gata1a*. Therefore, future work in this area may identify novel therapeutic targets for various disorders. For example, mutation of mevalonate kinase causes mevalonate kinase deficiency (MVD) ^70, 71^. Mevalonate kinase is central to the cholesterol synthesis pathway and converts mevalonate to 5’phosphomevalonate. 5’phosphomevalonate is the substrate for future enzymatic reactions that culminate with the creation of cholesterol and isoprenoids and therefore mutations in this kinase disrupt the synthesis of both cholesterol and isoprenoids. Patients with MVD exhibit with hematological deficiencies and extramedullary hematopoiesis ^72^, but the mechanisms underlying these phenotypes are not fully characterized. However, mutations in mevalonate kinase are likely to be recapitulated in zebrafish with mutations in *hmgcs1* or *hmgcrb*. Therefore, our system has the potential to understand the mechanisms governing *GATA1* expression, a central transcriptional regulator of primitive hematopoiesis.

## Supporting information

Virtual Summary

## Acknowledgements

These studies were supported by NIMHD Grant no.2G12MD007592 to University of Texas El Paso, NIGMS linked awards RL5GM118969, TL4GM118971, NIGMS grant no. R25GM069621-11, and UL1GM118970 to the University of Texas El Paso, and NINDS Grant no. 1K01NS099153-01A1 to AMQ. The *hmgcs1*^Vu57^ allele was kindly provided by Dr. Bruce Appel from the University of Colorado Anschutz Medical Campus. The *Tg*(*gata1a*:dsRed) fish were provided by Dr. Leonard Zon from Harvard Medical School.

## Author Contributions

JAH and AMQ synthesized the hypothesis, performed *in situ* hybridization, genotyping, data analysis, statistical analysis, cell counts, wrote the manuscript, and performed study design. JAH and VLC performed QPCR. AMQ and VLC performed morpholino injections and hemoglobin quantification in drug treatment assays. VLC performed *o*-dianisidine staining, imaged, and contributed to data analysis. NRN and JAH performed genotyping and imaging. JAH and LPM performed hemoglobin quantification in fish harboring the Vu57 allele.

