## Supplemental Data for "Mutations in the zebrafish *hmgcs1* gene reveal a novel function for isoprenoids during red blood cell development"

**Supplemental Figure 1**

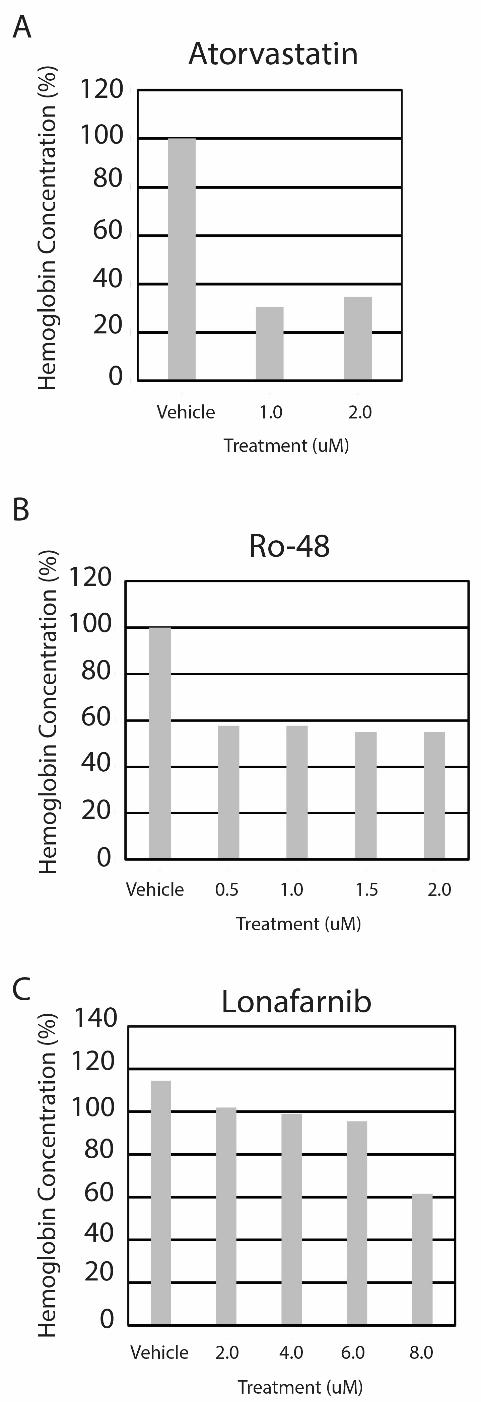

**Supplemental Figure 2**

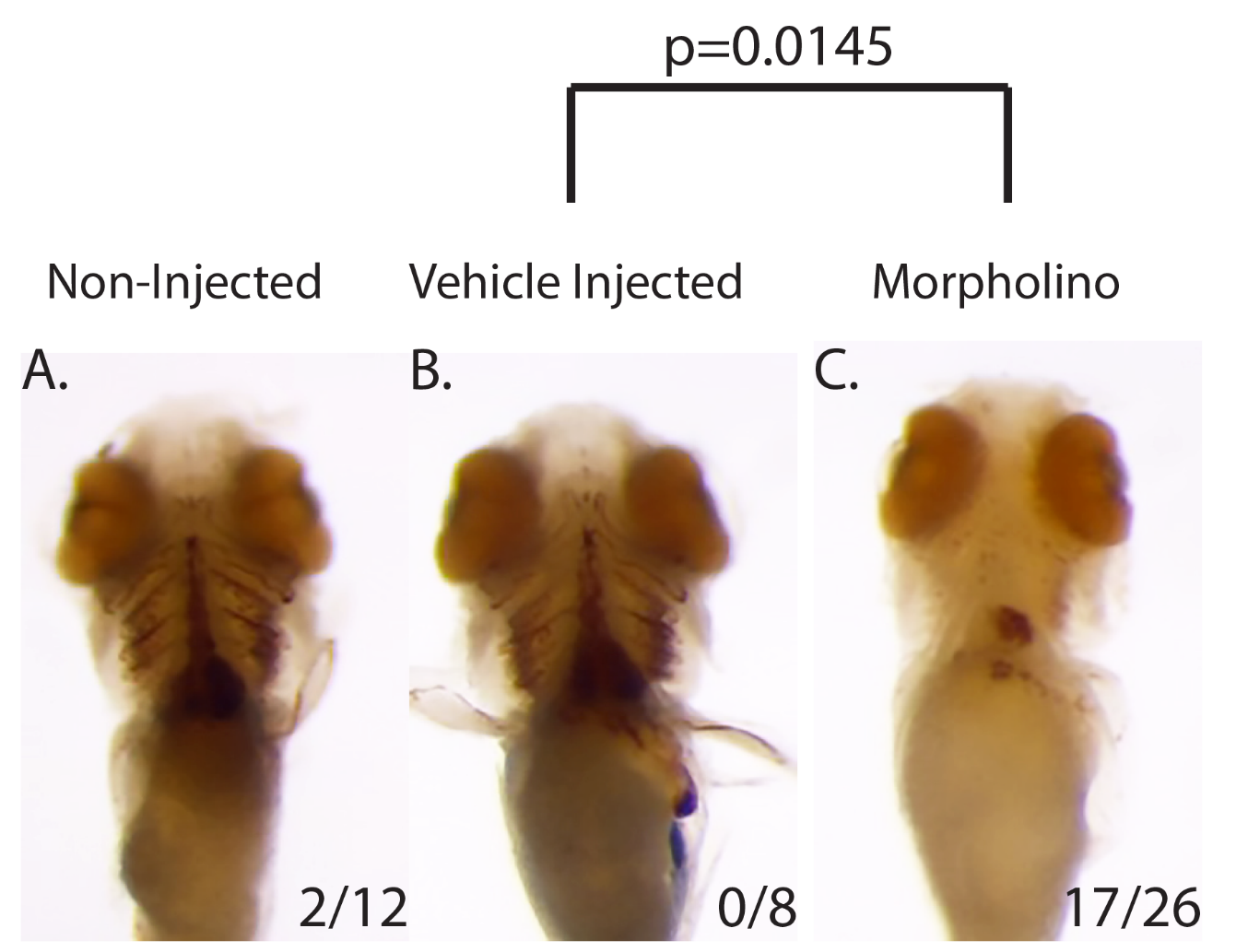

Supplemental Figure Legends

**Supplemental Figure 1:** A-C. The concentration of hemoglobin was measured in embryos treated with atorvastatin gradient (ATOR), Ro 48 8071 gradient (Ro-48) to inhibit cholesterol, l until 4 days post fertilization.

**Supplemental Figure 2:** A. Antisense *hmgcs1* morpholinos were injected (0.025mM) at the single cell stage and the number of RBCs were observed with *o*-dianisidine at 4 days post fertilization.
