## Supplementary figures and images for "Mutations in the zebrafish *hmgcs1* gene reveal a novel function for isoprenoids during red blood cell development"

### Virtual Summary

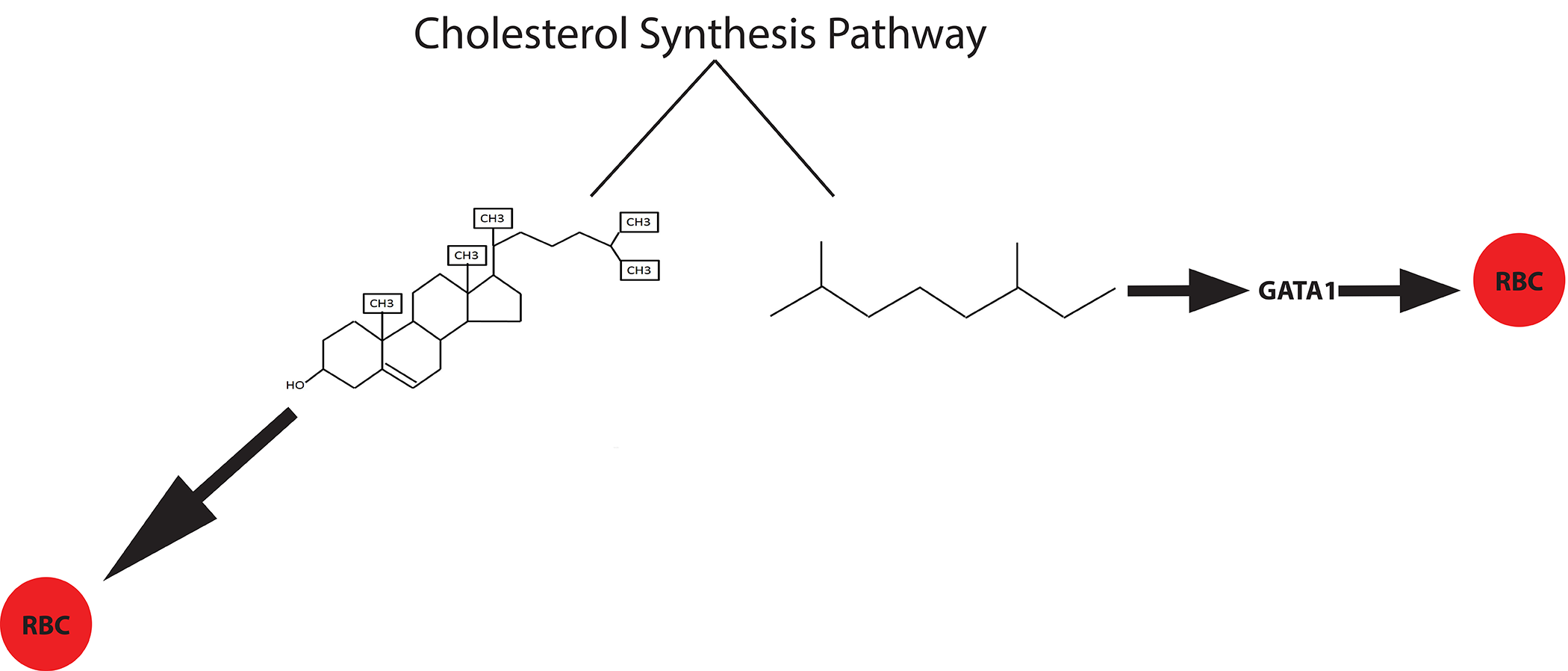
